# “Integrative Genomic Analysis for the Bioprospection of Regulators and Accessory Enzymes Associated with Cellulose Degradation in a Filamentous Fungus (*Trichoderma harzianum*)”

**DOI:** 10.1101/731323

**Authors:** Jaire A. Ferreira Filho, Maria Augusta C. Horta, Clelton A. dos Santos, Deborah A. Almeida, Natália F. Murad, Juliano S. Mendes, Danilo A. Sforça, Claudio Benício C. Silva, Aline Crucello, Anete P. de Souza

## Abstract

**Background:** Unveiling fungal genome structure and function reveals the potential biotechnological use of fungi. *Trichoderma harzianum* is a powerful CAZyme-producing fungus. We studied the genomic regions in *T. harzianum* IOC3844 containing CAZyme genes, transcription factors and transporters.

**Results:** We used bioinformatics tools to mine the *T. harzianum* genome for potential genomics, transcriptomics, and exoproteomics data and coexpression networks. The DNA was sequenced by PacBio SMRT technology for multi-omics data analysis and integration. In total, 1676 genes were annotated in the genomic regions analyzed; 222 were identified as CAZymes in *T. harzianum* IOC3844. When comparing transcriptome data under cellulose or glucose conditions, 114 genes were differentially expressed in cellulose, with 51 CAZymes. CLR2, a transcription factor physically and phylogenetically conserved in *T. harzianum* spp., was differentially expressed under cellulose conditions. The genes induced/repressed under cellulose conditions included those important for plant biomass degradation, including CIP2 of the CE15 family and a copper-dependent LPMO of the AA9 family.

**Conclusions:** Our results provide new insights into the relationship between genomic organization and hydrolytic enzyme expression and regulation in *T. harzianum* IOC3844. Our results can improve plant biomass degradation, which is fundamental for developing more efficient strains and/or enzymatic cocktails for the production of hydrolytic enzymes.

## Background

*Trichoderma harzianum* is a common fungal species in soil and is used as a biological control in a variety of phytopathogenic fungi [1]. However, the use of lignocellulosic biomass degradation is still poorly explored when compared to that of other cellulolytic fungi. Due to the high cellulolytic activity of some strains, *T. harzianum* has shown considerable potential for application in plant biomass hydrolysis [2–4]. *T. harzianum* strains have potential for the production of an enzymatic/protein arsenal necessary for the complete hydrolysis of cellulosic compounds in fermentable sugars [5–10].

Currently, the most-studied and widely used industrial-scale enzymes are produced by the fungus *T. reesei* and species from the *Aspergillus* genus. These organisms are the source of the majority of enzymes that make up enzymatic cocktails that are available on the market [11]. *T. reesei* is a widely studied fungus and is found in several works in genomics, transcriptomics, proteomics and metabolic engineering [12–16]. Thus, increasing the number of biotechnological studies related to this bioprocess for *T. harzianum* is necessary.

The three main groups involved in the hydrolysis of cellulose (CEL) are cellobiohydrolases, endo-β-1,4-glucanases and β-glucosidases. In addition, accessory enzymes such as copper-dependent lytic polysaccharide mono-oxygenases (LPMOs), cellulose-induced protein 1 and 2 (CIP1 and CIP2) and swollenin also participate in this process [17–20].

One of the great challenges in understanding the molecular mechanism of biomass degradation is how the transcription factors (TFs) related to this system act. Several fungal TFs have been identified as related to the degradation of plant biomass, many of which belong to the binuclear zinc family [21]. Many TFs have been described as being directly involved in the regulation of plant biomass [22]. This number has been expanding rapidly in recent years, mainly due to the increase in the sequencing scale of whole genomes and the exponential increase in bioinformatics tools for analysis, which produce massive amounts of information, and in the number of genes identified [22, 23].

The purpose of the present study was to analyze genomic regions with CAZyme genes using a bacterial artificial chromosome (BAC) library that we built [24] and to integrate these data with RNA-seq, secretome data and coregulation networks. We sequenced a massive amount of DNA and used it to integrate genomic data (genomic regions containing CAZymes), expression patterns (the transcriptome under degradation conditions), proteins (the secretome by mass spectrometry) and systems biology (with gene regulatory networks) to obtain a broad and precise overview of the CEL degradation pathways. Based on our study, we characterized the main genes, accessory enzymes and regions involved in the degradation and regulation process of hydrolytic enzymes. In addition, we analyzed the regulator cellulose degradation regulator 2 (CLR2) found in a cluster with other important enzymes. These results will be important for further studies of regulation and gene silencing.

## Results

### Genomic regions of T. harzianum IOC3844

In this study, a library of large genomic regions was used as a resource to search for genes of interest and to thoroughly study the genomic structure of *T. harzianum* IOC3844 (ThIOC3844) (accession numbers MK861589-MK861650 - Supplementary Table S1 and Fig. S1). Screening for genes of interest resulted in a total of 62 regions that contained CAZymes genes related to the degradation of plant biomass in the ThIOC3844 genome. Sequencing of these regions generated a total of 5 Mb of the estimated 40 Mb genome (Supplementary Table S2 and S3). These regions ranged in size from 43 to 152 kb, enabling the prediction and annotation of 1676 gene models for this strain (Supplementary Table S4). The average number of genes per region was 26 (Supplementary Table S1).

The genome of *T. reesei* QM6a (PRJNA325840) was used to analyze the distribution of genes in ThIOC3844. This genome, which is composed of seven chromosomes with a total size of 34 Mb, was divided into 38 intervals (1 Mb) (Fig. 1). It was possible to observe CAZyme genes annotated in ThIOC3844 distributed throughout the whole genome. Only four intervals had no CAZyme genes, and when all the genes in the genomic regions of ThIOC3844 were mapped, genes were found in all intervals.

**Figure 1.**
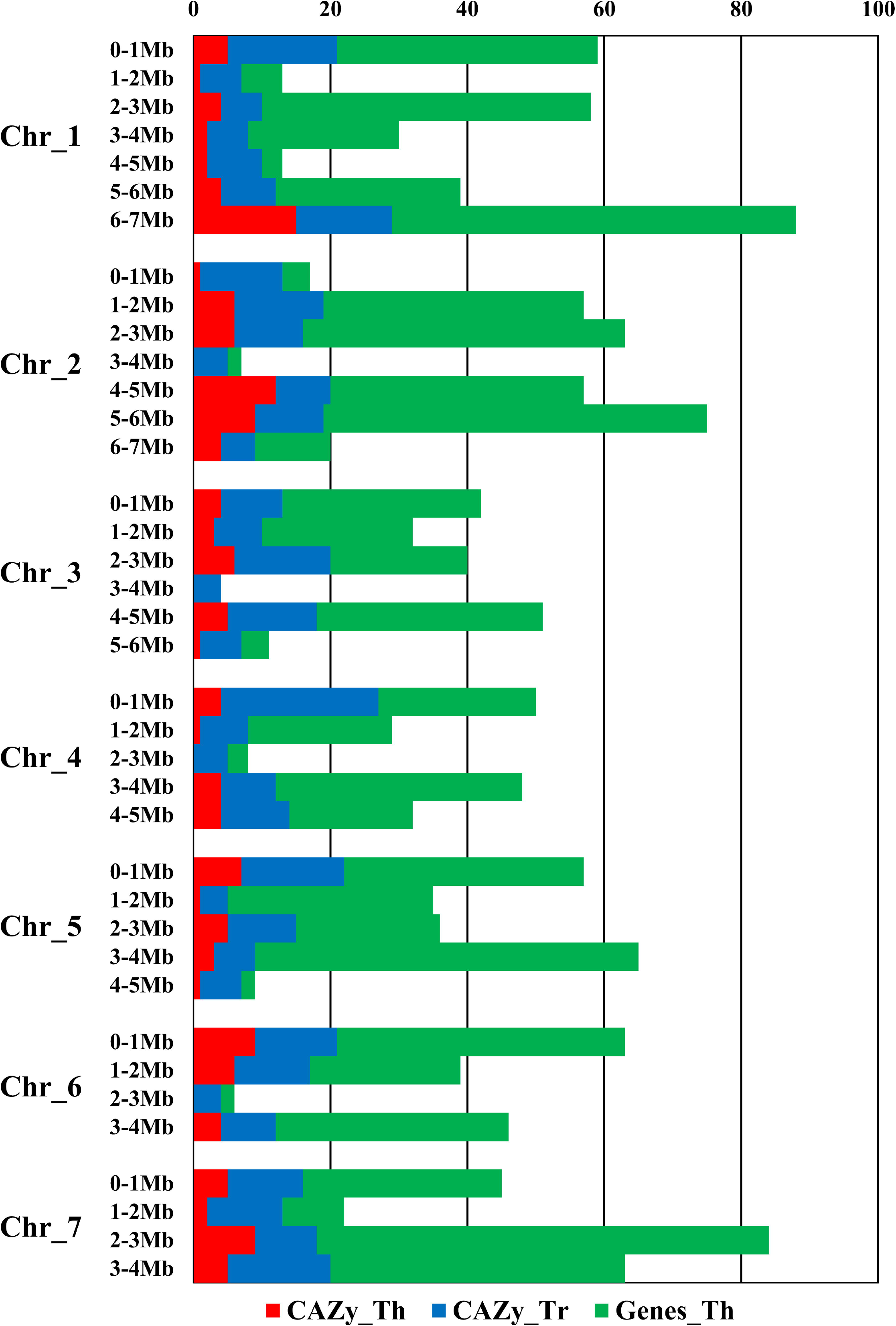
Distribution of the *T. harzianum* IOC3844 genes on the 1 Mb intervals of the seven chromosomes of *T. reesei* QM6a. CAZyme genes of *T. harzianum* IOC3844 are in red, CAZymes genes of *T. reesei* are in blue, and all genes of *T. harzianum* IOC3844 are in green. Th: *T. harzianum* IOC3844; Tr: *T. reesei*QM6a.

The genes were functionally annotated for the main gene ontologies: biological processes, cellular components and molecular functions (Fig. 2a and Supplementary Fig. S2). We found 209 sequences of hydrolytic activity, 139 related to transport proteins and 85 sequences involved in regulation of gene expression (possible TFs). In addition, a specific annotation was made for genes identified as enzymes, where hydrolases (40%), oxidoreductases (25%), transferases (22%), lyases (6%), ligases (4%) and isomerases (3%) (Figure 2b) were found. We also identified genes directly related to the degradation of CEL and hemicellulose, with action of α-L-arabinofuranosidase (EC 3.2.1.55), endo-1,4-β-xylanases (EC 3.2.1.8), cellobiohydrolases (3.2.1.91), endo-β-1,4-glucanase (EC 3.2.1.4) and β-glucosidase (EC 3.2.1.21) (Fig. 2c and Supplementary Table S5).

**Figure 2.**
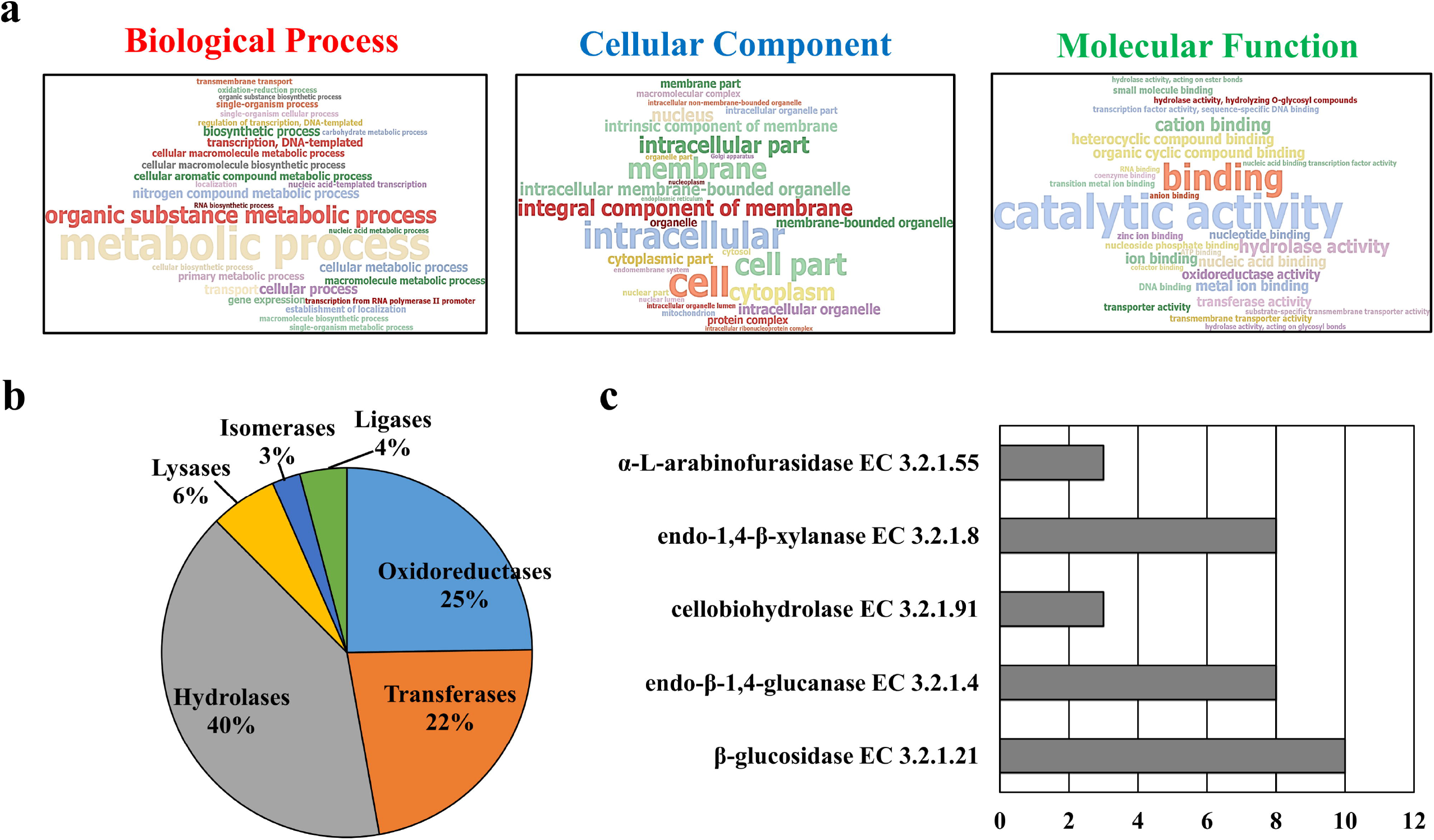
Functional annotation of the genes predicted in the genomic regions of *T. harzianum* IOC3844. Annotation of genes for gene ontologies for biological processes, cellular components and molecular functions. (a) Distribution of enzymes annotated according to enzyme commission (b) and major enzyme commission (EC) related to cellulose and hemicellulose degradation (c).

A total of 1676 genes were predicted. Of these, 222 were annotated as CAZymes in ThIOC3844, including 45% of GHs, 23% of GTs, 10% of CEs, 8% of AAs and 14% of CBMs (Fig. 3 and Supplementary Table S6). The GH class presented with the highest number of families, including GH2 (3 genes), GH7 (1 gene), GH3 (9 genes), GH5 (6 genes), GH12 (1 gene), GH18 (4 genes) and GH62 (1 gene).

**Figure 3.**
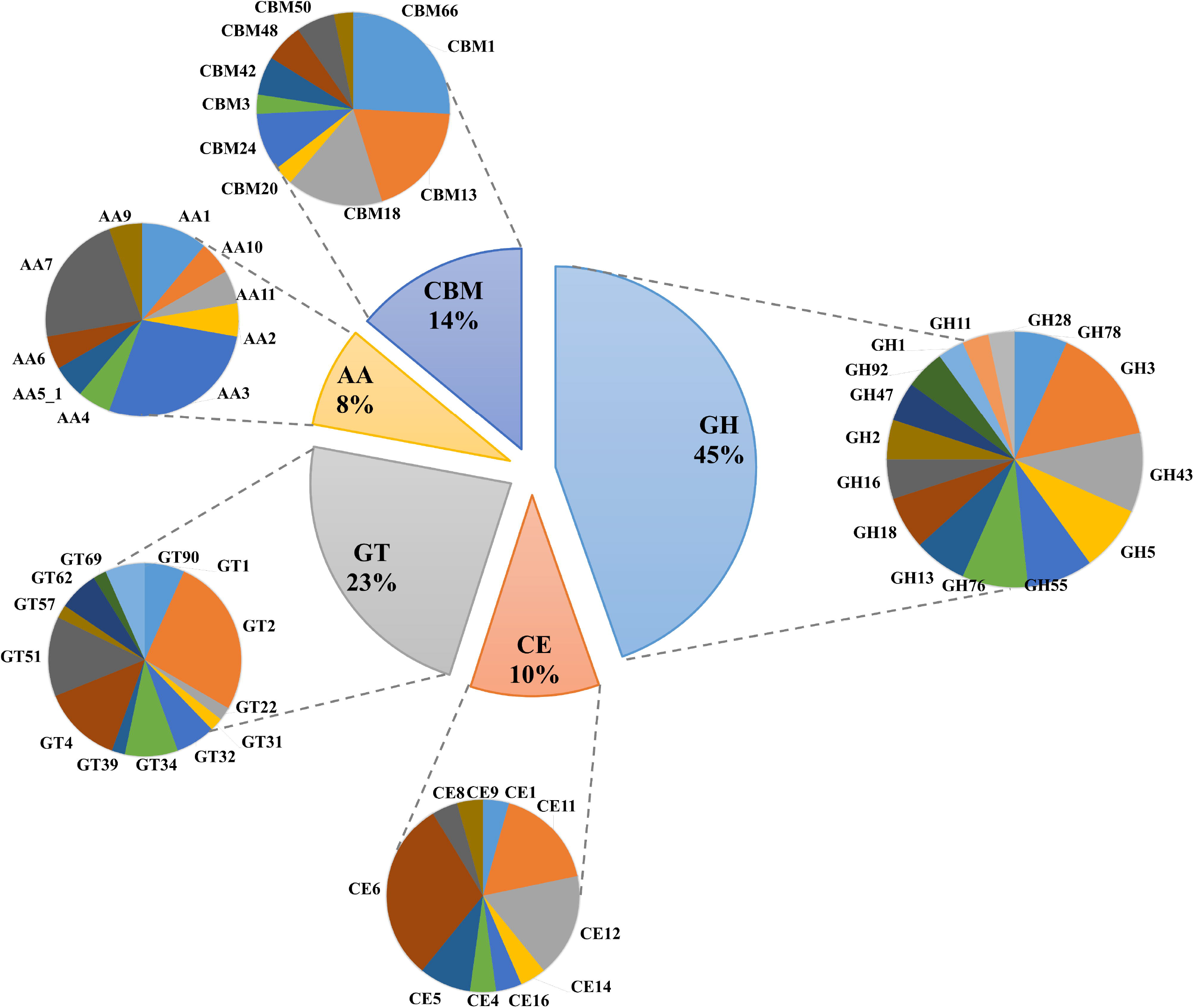
CAZy classification of genes annotated in the genomic regions of *T. harzianum* IOC3844. GH: glycoside hydrolases; GT: glycosyl transferases; PLs: polysaccharide lyases; CEs: carbohydrate esterases; AA: auxiliary activities; CBM: carbohydrate-binding modules.

### Genomic comparison

For this analysis, we compared the genomic regions of ThIOC3844 against the entire genome of different strains and species of the genus *Trichoderma*. Genomic comparison of the sequenced regions of ThIOC3844 with two other strains of the same species *(T. harzianum* B97 – ThB97 and *T. harzianum* – T6766) showed a higher similarity to ThB97 (99.25%) than ThT6766 (91.61%). For the *T. atroviride* IMI206040 genome (TaIMI206040), the similarity to ThIOC3844 was 85.09%. For *T. virens* Gv29-8 (TvGv29-8), the similarity was 86.55%, and for *T. reesei* QM6a (TrQM6a), the similarity was 85.11%.

When we compared syntenic genes between groups of genes, a greater difference between *T. harzianum* and *T. atroviride* and *T. reesei* was observed. The *T. harzianum* TR274 (ThTR274) strain presented the same gene profile of genomic organization as that found in ThIOC3844. In TaIMI206040, four genes (GH4, transporter and two GH26) from the cluster were not found; for TvGv29-8, two genes were not found (GH1 and GH4). For *T. reesei* QM6a, three genes (GH4 and two GH26) were not found; in addition, the translocation of genes (MFS x GH2 and TF2 x CLR2) was found. The genes for the transcription factor CLR2, putative transcription factor TF2 and MFS (major facilitator superfamily permease) were maintained in all species analyzed. This result suggests a potential association between the regulation and expression of these genes (Fig. 4).

**Figure 4.**
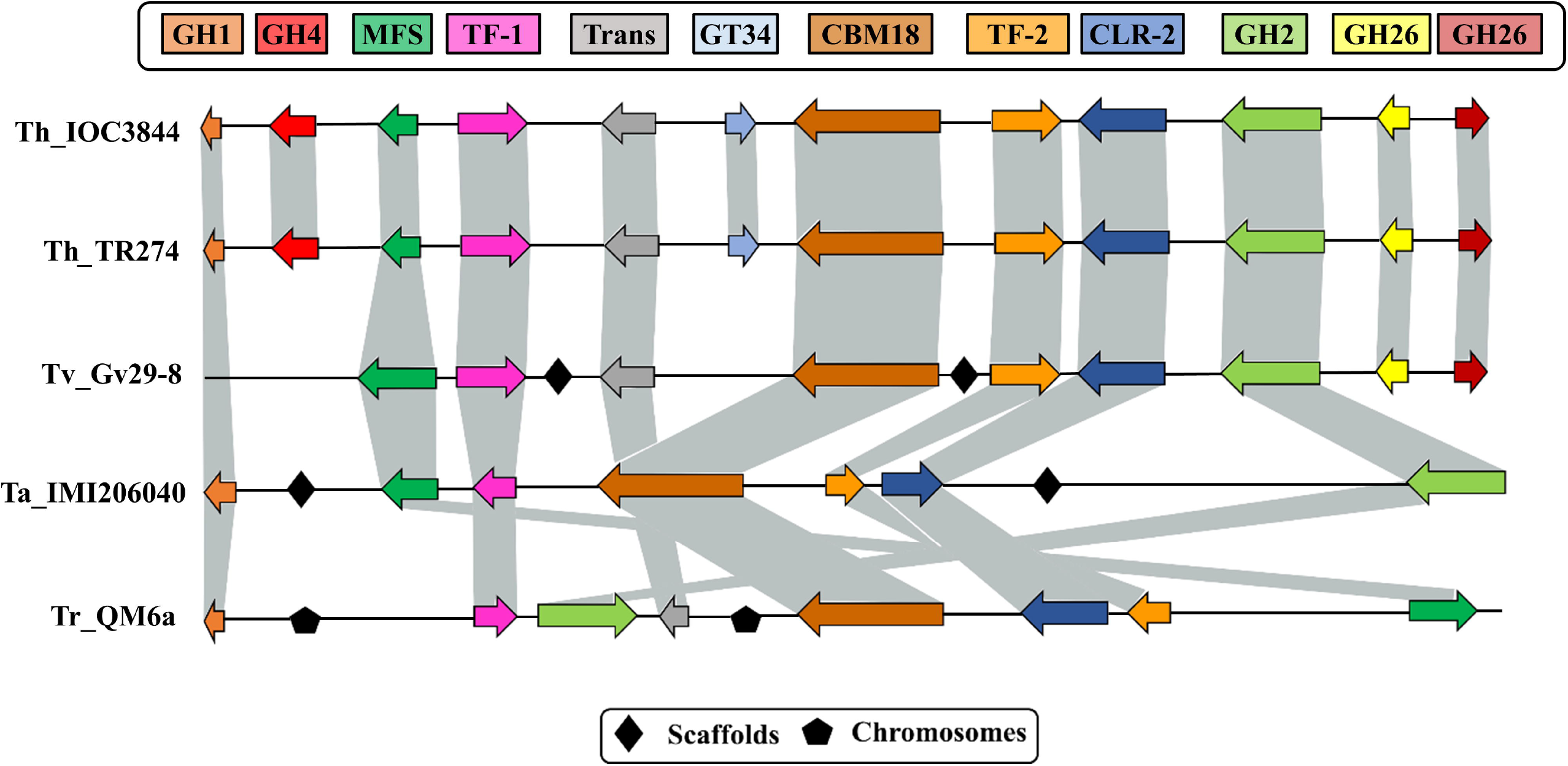
Comparison between the gene clusters of *T. harzianum* IOC3844 and those of other species of the genus *Trichoderma* spp. GH1: glycoside hydrolase 1; GH4: glycoside hydrolase 4; MFS: major facilitator superfamily permease; Trans: putative transporter; TF-1: putative transcription factor 1; GT38: glycosyl transferases 4; CBM18: carbohydrate-binding modules 18; TF-2: putative transcription factor 2; CLR2: cellulose regulator 2; GH2: glycoside hydrolase 2; GH26: glycoside hydrolase 26; Th: *T. harzianum;* Tv: *T. virens;* Ta: *T. atroviride;* Tr: *T. virens*.

### Expression by RNA-Seq and secreted proteins

All genes predicted in the genomic regions were analyzed according to expression data by RNA-Seq (under CEL and GLU degradation conditions) (Supplementary Table S7) and secreted proteins identified by mass spectrometry (LC-MS/MS). We found 114 genes with differential expression under CEL degradation conditions when compared to GLU degradation conditions; among them, 51 were classified as CAZymes, such as beta-glucosidase of the GH1 family (1.8-fold change - FC), LPMOs of the AA9 family (FC 5.0) and hypothetical protein with domain CBM1 (FC 3.7). In addition, two differentially expressed TFs were identified, CLR2 (FC 1.6) and unidentified transcriptional regulator of zing finger – Zn2Cys6 (FC 2.3). Six transport proteins were also found (iron permease, MFS hexose transporter, siderophore transporter, ammonium permease, sugar transporter and siderophore iron transporter).

Among the genes annotated as CAZymes in ThIOC3844, 31 were found in the secretome of ThIOC3844 under CEL conditions, and the main families were GH3, GH12, CBM1, AA9, GH6/CBM1, GH45/CBM1, GH62 and GH5. In this analysis, we also used the level of expression of the secreted genes. The gene with the highest TPM index (1567.4 TPM) is a cellobiohydrolase (EC 3.2.1.91) of the GH6 family. However, our results indicate that genes with low expression levels are also important secreted enzymes (Table 1).

**Table 1.** Proteins identified in genomic and in the *T. harzianum* IOC3844 secretome under cellulose growth conditions.

| **IDs\*** | **Protein name** | **Secretome/UniPro**<br>**t ID** | **CAZy**<br>**family** | **CEL**<br>**(TPM)** | **GLI**<br>**(TPM)** |
| --- | --- | --- | --- | --- | --- |
| 1010 | Hypothetical protein | A0A0G0ALT6 | GH28 | 14.2 | 6.2 |
| 1043 | Cellulosome enzyme | A0A0G0A296 | GH30 | 35.6 | 11.5 |
| 1054 | Glycosyl hydrolase<br>10 | A0A0F9X8A4 | GH10 | 14.8 | 4.1 |
| 1075 | Glycosyl hydrolase<br>64 | A0A0F9ZIR5 | GH64 | 824.3 | 262.4 |
| 1095 | Glycosyl hydrolase<br>18 | A0A0F9ZHI0 | GH18 | 83.2 | 47.9 |
| 11 | Mutanase | A0A0F9XN06 | CBM24 | 2741.6 | 1452.9 |
| 1133 | Glycosyl hydrolase<br>12 | A0A0F9Y2E9 | GH12 | 1579.8 | 308.2 |
| 1150 | Glycosyl hydrolase<br>47 | A0A0F9WYR7 | GH47 | 83.9 | 74.6 |
| 1217 | Beta-mannosidase | A0A0F9ZDV4 | GH2 | 117.9 | 124.2 |
| 126 | Glycosyl hydrolase<br>76 | A0A0F9X1Q3 | GH76 | 616.7 | 375.4 |
| 1318 | Beta-xylosidase | A0A0G0A408 | GH3 | 172.3 | 125.2 |
| 1439 | Alpha-L-arabinofuranosidase<br>B | A0A0G0A4Q2 | CBM42 | 450.4 | 343.5 |
| 1440 | Glycosyl hydrolase 3 | A0A0F9XRC5 | GH3 | 245.8 | 107.4 |
| 1498 | WSC domain-<br>containing | A0A0F9ZXC9 | AA5_1 | 342.5 | 339.0 |
| 44 | Beta-1,3- | A0A0F9ZKA8 | GH72 | 2431.7 | 3210.5 |

| glucanosyltransferase |  |  |  |  |  |
| --- | --- | --- | --- | --- | --- |
| 441 | Alpha-glucosidase | A0A0G0AG54 | GH31 | 2121.6 | 1655.2 |
| 559 | Alpha-1,2-mannosidase | A0A0G0ABI9 | GH92 | 226.9 | 153.4 |
| 666 | Glycosyl hydrolase 3 | A0A0F9XQT4 | GH3 | 77.2 | 43.0 |
| 667 | Hypothetical protein | A0A0G0AME2 | CBM1 | 874.9 | 142.8 |
| 668 | Glycosyl hydrolase 61 | A0A0F9XMI8 | AA9 | 3109.7 | 625.1 |
| 669 | Glycosyl hydrolase 16 | A0A0F9XP75 | CBM13 | 16.4 | 3.7 |
| 671 | Cytochrome P450 monooxygenase | A0A0G0A4Z5 | GT4 | 1569.5 | 1595.3 |
| 681 | Glycosyl hydrolase 11 | A0A0F9Y0Y9 | GH11/CBM1 | 4206.8 | 1316.1 |
| 741 | Endo-N-acetyl-beta-D-glucosaminidase | A0A0F9ZHA7 | GH18 | 3971.9 | 2328.0 |
| 759 | Hypothetical protein | A0A0F9ZJ74 | GH20 | 1184.8 | 1507.7 |
| 813 | Catalase peroxidase | A0A0F9X3Z8 | AA2 | 2677.7 | 2473.4 |
| 82 | Glycosyl hydrolase 6 | A0A0G0AEM7 | GH6/CBM1 | 5843.5 | 1567.4 |
| 842 | Hypothetical protein | A0A0F9XY55 | GH45/CBM1 | 41.3 | 15.3 |
| 9 | Glycosyl hydrolase 62 | A0A0F9X8Z0 | GH62 | 353.9 | 103.4 |
| 913 | Isoamyl alcohol oxidase | A0A0F9XC99 | AA7 | 39.3 | 13.2 |
| 918 | Hypothetical protein | A0A0F9XG06 | GH5_5 | 870.8 | 750.9 |

\*The annotated genes IDs can be found in Supplementary Table S4

### CLR2 transcription factor

The phylogenic analysis of the CLR2 factor showed a clear separation of this TF in relation to Basidiomycetes and Ascomycetes (Fig. 5a and Supplementary Table S8). However, even within these groups, considerable phylogenetic diversity was observed among the species of analyzed fungi with a variety of clades within the same group. Different strains of *T. harzianum* grouped in a single clade with proximity to *T. reesei* and *T. atroviride* species. Our results show a wide range of functional variety for CLR2, which may indicate different types of performance between species.

**Figure 5.**
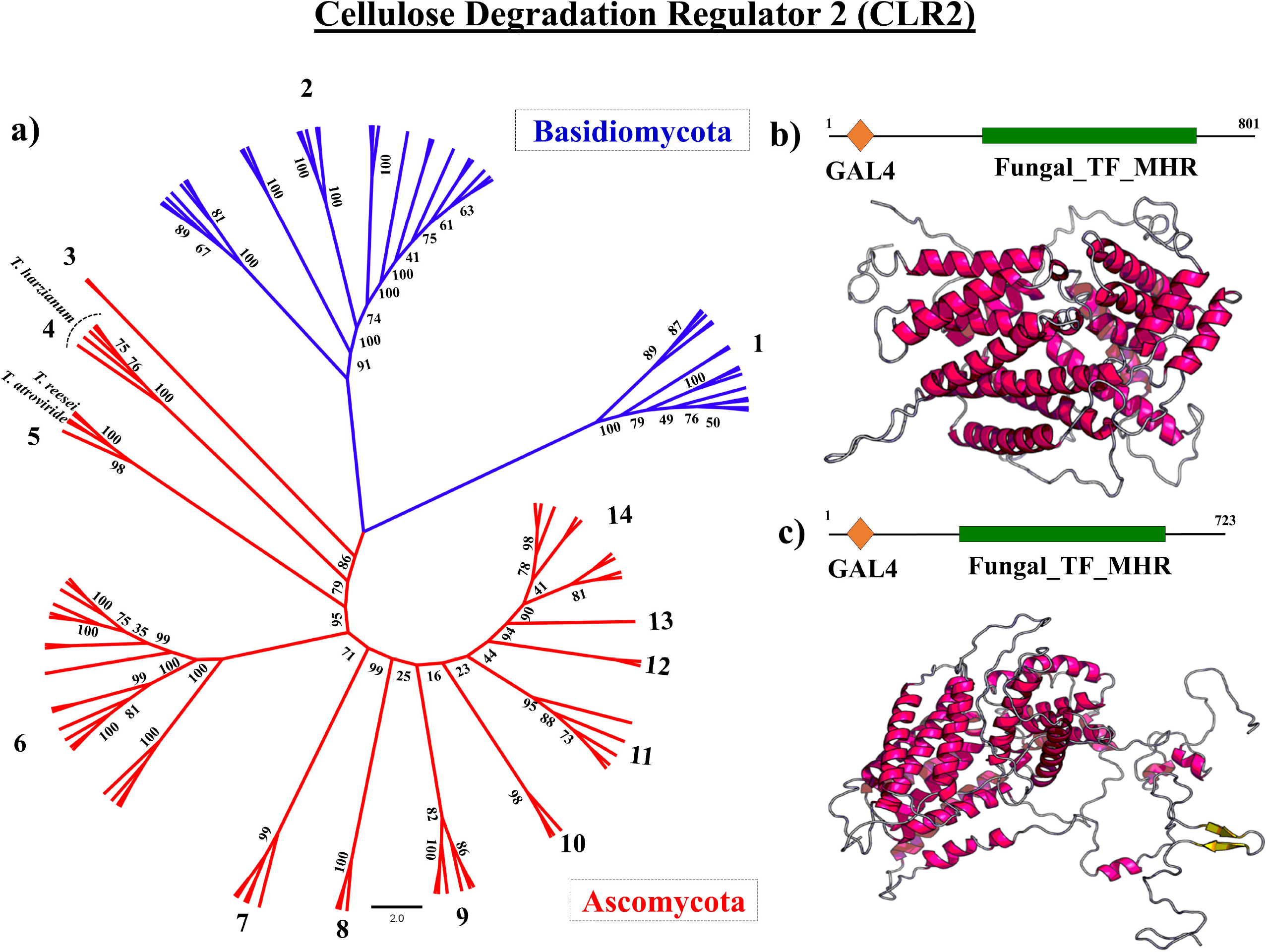
Molecular phylogeny of the CLR2 transcription factor in Ascomycota and Basidiomycota (a); *in silico* protein modeling for CLR2 in *T. harzianum* IOC3844 (b) and *T. reesei* QM6a (c).

A structural modeling analysis for the CLR2 protein of ThIOC3844 was performed using *T. reesei* as a comparator. For both proteins, the best template was <u>6F07</u> (*Saccharomyces cerevisiae*), with e-values of 4.07e^-06^ and 6.62e^-06^ for ThIOC3844 (Figure 5b) and *T. reesei* (Figure 5c), respectively. Prediction of 1 and 3 protein domains was made for ThIOC3844 and *T. reesei*, respectively. For ThIOC3844, 59% of the residues were already modeled, and for *T. reesei*, it was possible to model 83%. For ThIOC3844, the secondary structure prediction was 46% H (helix), 0% E (beta-sheet) and 53% C (loop), and for solvent access, it was 56% E (exposed), 19% M (medium) and 23% B (buried).

A coregulation network of genes directly related to the CLR2 regulator was constructed, searching for insights about other important proteins in the process of cellulase expression. We identified 36 genes directly linked to CLR2, of which 21 genes were annotated as hypothetical proteins. In addition, we found that genes with known annotations were related to the process of gene expression, including genes annotated as initiation factors, kinases and helicases (Fig. 6a and Supplementary Table S9).

**Figure 6.**
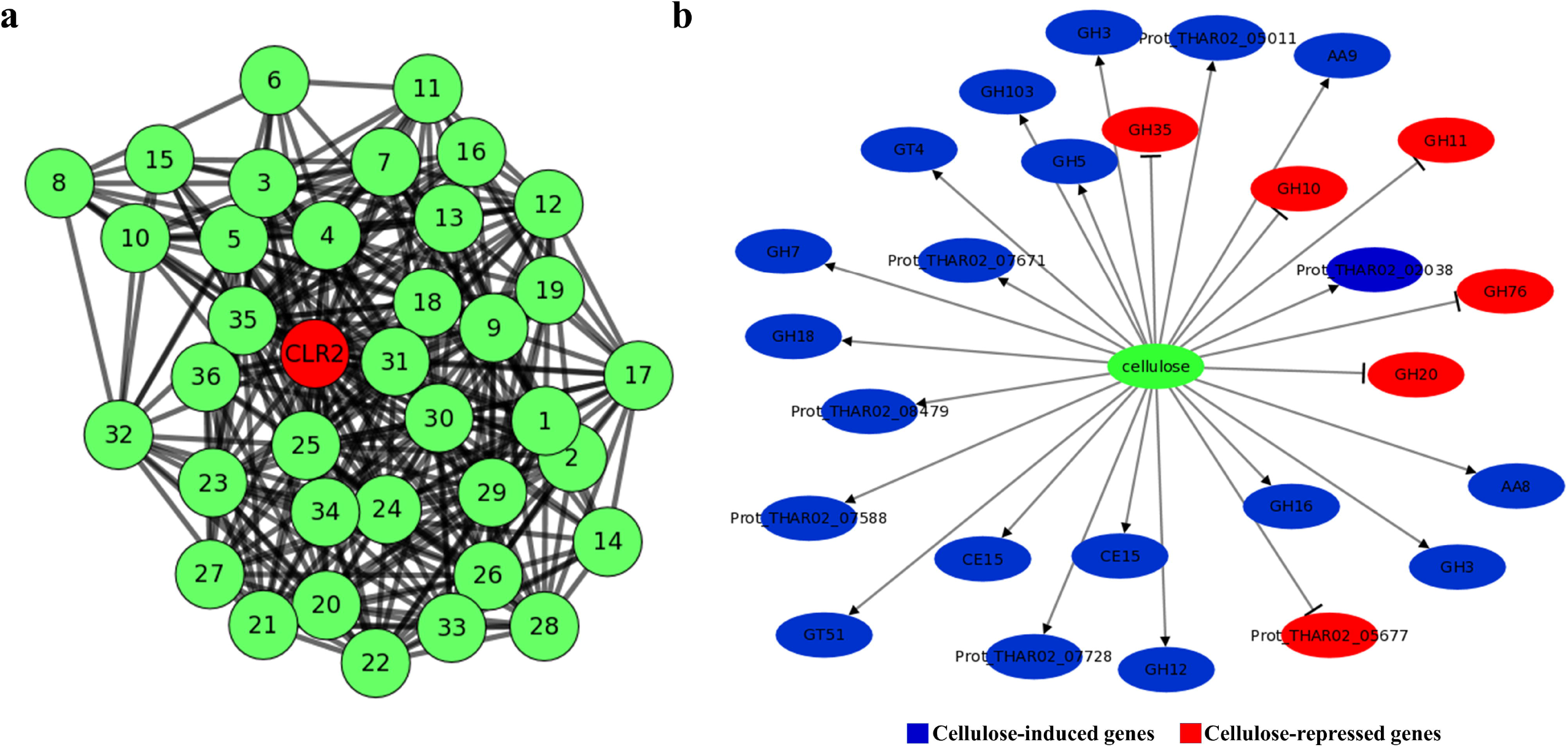
Subnetwork of CLR2 transcription factors and related genes (a) and network of induced (blue) and repressed (red) genes under cellulose conditions (b). CLR2: cellulose regulator 2; GH: glycoside hydrolases; GT: glycosyl transferases; AA: auxiliary activities.

### Network of induced/repressed genes in cellulose

Using the gene expression data of the secreted proteins, a Bayesian network of induced/repressed genes was constructed based on the CEL growth conditions for *T. harzianum* IOC3844 (Fig. 6b). The major genes that were induced under this condition belong to the GH7 (exoglucanase), GH5 (endo-β-1,4-glucanase), GH3 (β-glucosidase), GH12 (murein transglycosylase), CE15 (CIP2), AA9 (LPMO) and AA8 (hypothetical protein) families. In addition, seven genes that were not classified as CAZymes were also induced under CEL conditions. The families of repressed genes were GH10 (glycoside hydrolase 10 family endo-1,4-β-xylanase), GH11 (glycoside hydrolase 11 family endo-1,4-β-xylanase), GH76 (alcohol dehydrogenase 1), GH20 (β-N-acetylhexosaminidase) and GH35 (glycoside hydrolase 35).

## Discussion

In the present study, an integrative multi-omics approach was used to mine CAZyme-rich regions of ThIOC3884. BAC clones were selected, sequenced and used in comparative analyses focusing on the expression profile via RNA-Seq and the exoproteome under different fungal growth conditions, enabling the discovery of important gene/proteins related to plant biomass degradation (Supplementary Fig. S3).

The vast majority of important enzymes for the degradation of plant biomass are already known [25–27]. The current challenge is how enzymes are regulated and the genetic mechanism of their activation. Thus, many works with cellulolytic fungi have focused on TFs, accessory enzymes, transporters and the way the type of biomass affects the process of regulating the cellulases and hemicellulases [22, 28–30]. Other studies have already shown the potential of *T. harzianum* for the degradation of plant biomass. This is the first work that integrates results from different biotechnology approaches and that focuses on the prediction of the most important enzymes and TFs used by *T. harzianum* IOC3844 to degrade CEL.

The molecular process of CEL degradation is extremely complex and involves hydrolytic enzymes acting on the extracellular medium, carrier proteins and TFs (Figure 7). For *T. harzianum* and *T. reesei*, the major CAZy families related to CEL degradation were identified in the genome (GH1, GH3, GH6, GH7, GH12, GH45 and AA9) [7], and many of the cellulases have already had their three-dimensional structure solved; however, many key proteins in this process are not well known as transporter TFs related to the regulation of these enzymes.

**Figure 7.**
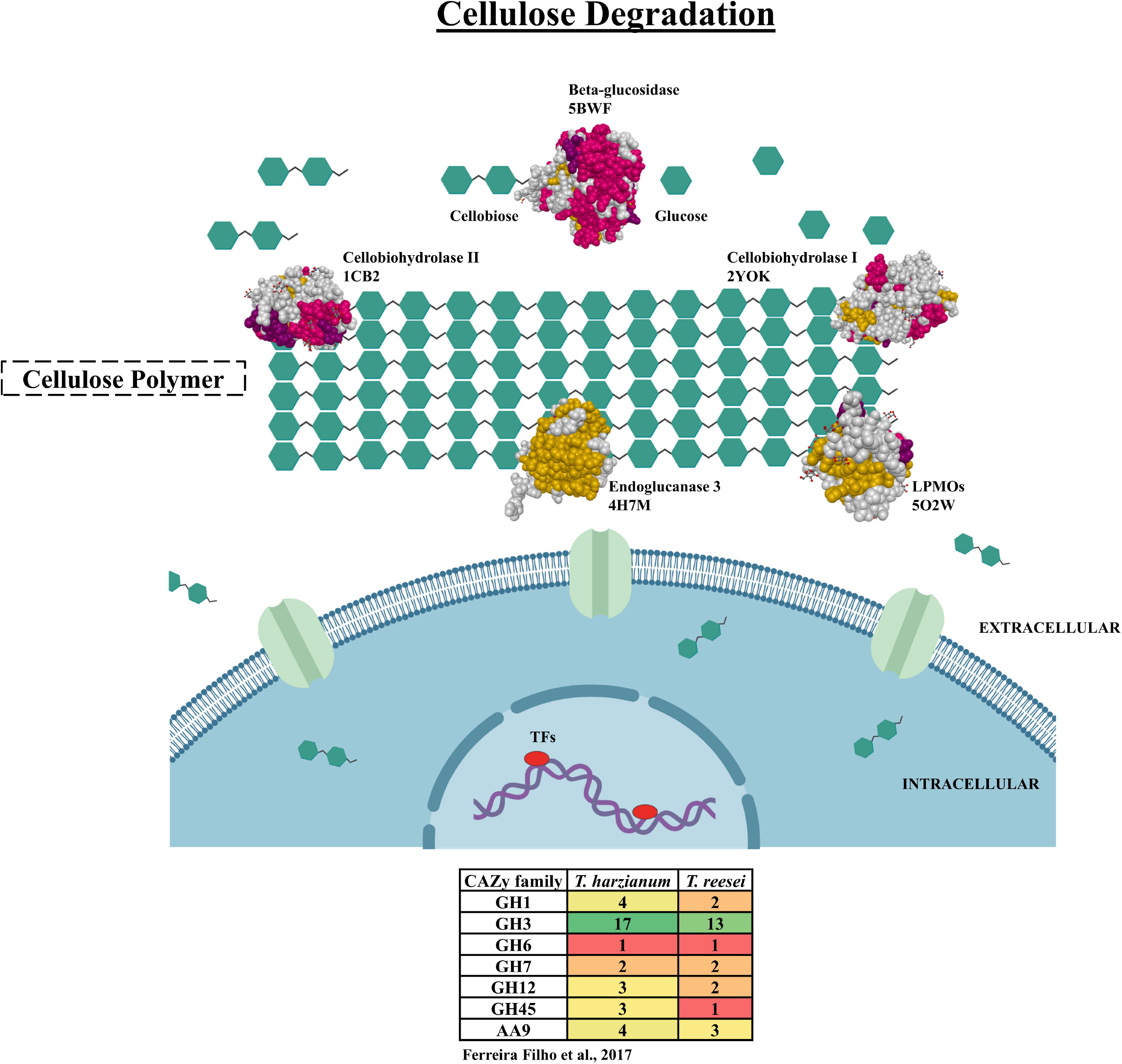
Molecular scheme of the enzymatic model in the degradation of cellulose in *Trichoderma* spp. Enzymes and PDB code: beta-glucosidase (<u>5BWF</u>), cellobiohydrolase I (<u>2YOK</u>), cellobiohydrolase II (<u>1CB2</u>), endoglucanase 3 (<u>4H7M</u>), copper-dependent lytic polysaccharide mono-oxygenases (LPMOs) (<u>5O2W</u>).

The study of genomic regions is an important tool for providing a global view of the important genes and regulatory regions of a genome [24, 31]. The genomes of a few strains of *T. harzianum* are available [32, 33]. A complete genome draft sequenced in 1572 scaffolds is available for *T. harzianum* T6776 [32]; however, little is known about the ThIOC3844 genome, and as it is a strain with potential for hydrolytic enzymes, more genomic information regarding CAZyme sequences is needed. In this study, our strategy was to use large genomic regions and integrate these data with other genetic information.

A large number of fungal genomes have already been used as a platform to search for new genes related to the degradation of biomass, as is the case for *T. reesei* QM6a, which has a finalized genome divided into seven chromosomes [34]. Our study results with the genomic regions of ThIOC3844 showed a large number of enzymes classified as CAZymes, as well as TFs and transporters in clusters in the genome, which may be important for future studies of genetic modification of this lineage.

Analyzing the level of expression of certain genes under certain conditions is an important step in understanding how transcription is affected in a specific biological condition [17, 35]; however, there is not always a direct relationship between what is being highly expressed and the proteins that are important in the extracellular medium. Thus, in this work, in addition to studying the most expressed genes that we found in the genomic regions, we also searched for those with a confirmed presence in the fungus secretome CEL degradation conditions. Our results showed that CAZy families are key in the degradation of CEL, with a high level of expression and a positive presence as a secreted protein.

Genomic comparison is a powerful tool for understanding differences and evolutionary dynamics among related species [36–38]. Our data show a high similarity between different strains of *T. harzianum* (IOC3844, B97 and T6776), which indicates that differences in enzyme production and efficiency may be related more to gene regulation mechanisms than differences in the sequence itself. In addition, by synteny analysis, it was possible to observe a greater difference in relation to the genome of *T. reesei*, which can be explained by the loss of genes and genomic modifications carried out in lineages of this fungus to increase its productivities of enzymes related to plant biomass degradation [12, 39].

The CLR2 transcription factor was described as an important regulator in the expression of cellulases by *Neurospora crassa* [22]; however, its functional role is not yet clear for fungi of the genus *Trichoderma*, including *T. reesei* [14, 40]. In the genome of ThIOC3844, we found a cluster with the CLR2 TF in association with other putative transcription factors, CAZymes, transporters and MFS permease. The same behavior was found for the *T. reesei* CLR2 TF, which has physical proximity and coexpression with a sugar transporter [29, 41]. These results indicate that there may be a mechanism for the joint regulation and expression of this TF with transporters related to biomass degradation. Based on RNA-Seq data, we observed differential expression of CLR2 in the cellulose condition. In this way, we analyzed the coregulation network of the CLR2 regulator. The present study illuminates unclear areas of the genomic organization, expression and putative regulation of CLR2 in *T. harzianum*.

Coregulation networks provide insights into how genes correlate and interact with each other [35, 42, 43]. We identified 36 genes directly associated with the CLR2 regulatory factor; these genes may be important in the regulation process of this factor, which is linked to the expression of cellulases in other filamentous fungi. Techniques such as gene knockout can further validate the functional or synergistic importance of these genes with key TFs for the expression of genes related to degradation of plant biomass.

## Conclusions

Our results present an innovative approach in using different types of omics data to search for new important genes and genetic regulation mechanisms during the process of CEL degradation. We found several TFs, accessory enzymes and transporters in the genomic regions of ThIOC3844 that may be important for the expression/secretion of CAZyme genes. Among these, CLR2, CIP2 and LPMOs are promising candidates for further study. Our results indicate that the CRL2 regulator matches all the requirements for involvement in cellulose degradation by *T. harzianum*. In addition, through the approach of coregulation networks, it is possible to understand the relationship between genes and to find new targets for biochemical characterization. The results allowed the identification of important genetic regions, key genes and functional proteins, and this information can be used for the development and improvement of enzymatic hydrolysis technology for the bioethanol industry.

## Methods

### T. harzianum *strain and genomic resources*

*T. harzianum* IOC3844 (ThIOC3844) was obtained from the Brazilian Collection of Environment and Industrial Microorganisms (CBMAI). A library of BACs consisting of 5,760 clones previously constructed for this fungus strain [24] was used to search for genomic regions. The genomic sequences of *T. harzianum* T6776 (<u>PRJNA252551</u>), *T. reesei* QM6a (<u>PRJNA325840</u>), *T. atroviride* IMI206040 (<u>PRJNA19867</u>) and *T. virens* Gv29-8 (<u>PRJNA19983</u>) were used for comparison with ThIOC3844.

### *BAC library screening for gene selection in* T. harzianum *IOC3844*

We designed primers for 62 target CAZyme genes (Supplementary Table S1) using transcriptome data [3] to search for positive BAC clones that contain genes previously selected from the plate (with the complete BAC library comprising fifteen 384 plaques) and column pools (24 columns of each plate). The plate and column pools were amplified using the Illustra GenomiPhi HY DNA Amplification Kit (GE Healthcare Life Sciences, UK) following the manufacturer’s instructions. The screening reactions for the search for positive clones were performed via PCR using the CFX384 Touch Real-Time PCR Detection System (Bio-Rad).

### Single-molecule real-time (SMRT) sequencing and assembly

Libraries for sequencing were prepared according to the Pacific Biosciences (PacBio) protocol, and sequencing was performed at the Arizona Genomics Institute (AGI; Tucson, USA) using a Single-Molecule Real-Time (SMRT) DNA sequencing system available from PacBio. *De novo* assembly was performed with the PacBio Corrected Reads (PBcR) pipeline implemented as part of Wgs-assembler v8.3rc2 [44] and Celera Assembler [45]. The contigs obtained with the assemblers were subjected to error correction with pbalign (v0.2). The PacBio reads were aligned using the BLASR algorithm [46], and assembly polishing was performed with the Quiver tool (Supplementary Table S2 and S3) [47].

### Gene prediction and functional annotation

The FGENESH tool was used for initial gene prediction analysis [48], followed by manual correction with the *T. harzianum* T6776 and *T. reesei* QM6a gene models. Annotations of the ontologies were performed with Blast2GO [49]. InterPro protein domains were predicted using InterProScan (http://www.ebi.ac.uk/interpro/) [50]. Information derived from the CAZy database was downloaded for each CAZyme family (www.cazy.org). The protein sequences of *T. harzianum* IOC3844 were used as queries in basic local alignment search tool (BLASTp) searches against the locally built CAZyme BLAST database. Only BLAST matches showing an e-value less than 10^-11^, identity greater than 30% and queries covering greater than 70% of the sequence length were retained and classified according to the CAZyme catalytic group as glycoside hydrolases (GHs), glycosyl transferases (GTs), polysaccharide lyases (PLs), carbohydrate esterases (CEs), carbohydrate-binding modules (CBMs) or auxiliary activities (AAs).

### *Genomic comparison in* Trichoderma spp

The software used for alignment was Nucmer (-maxmatch), which is part of the software package MUMmmer 3.23 [51]. The delta-filter (-q), show-coords (-rcl), and DNADIFF (standard parameters) were used for filtering, obtaining the mapping coordinates and generating the statistical report in the alignment, respectively. SimpleSynteny software (https://www.dveltri.com/simplesynteny/) [52] was used to compare a cluster of 12 genes among different species of *Trichoderma* spp.

### Phylogenetic analysis and structure modeling of CLR2

The CLR2 sequences of ThIOC3844, *T. reesei* QM6a, *T. atroviride, T. virens* and other species of fungi were used as the basis for constructing the phylogenetic trees. These fungi were divided into Ascomycetes and Basidiomycetes. The sequences were aligned using ClustalW [53] and analyzed with Molecular Evolutionary Genetics Analysis (MEGA) software v7.0 (https://www.megasoftware.net/) [54]. The phylogenetic analyses were performed in MEGA7 using the maximum likelihood (ML) [55] method of inference based on the Jones-Taylor-Thornton (JTT) matrix-based model and 1000 bootstrap replicates [56] for each analysis. Pairwise deletion was employed to address alignment gaps and missing data. The trees were visualized and edited using the FigTree program (http://tree.bio.ed.ac.uk/software/figtree/). *In silico* modeling of the domain of CLR2 was performed using RaptorX protein structure prediction software (http://raptorx.uchicago.edu/) [57].

### RNA-Seq and exoproteome analysis

The expression levels of ThIOC3844 were analyzed using RNA-Seq data (<u>PRJNA336221</u>) obtained from a previous study in which the transcripts were obtained following growth of the fungus on two different carbon sources, CEL and GLU [35]. The reads from the RNA-Seq library were mapped against the ThIOC3844 genes using the CLC Genomics Workbench (https://www.qiagenbioinformatics.com/products/clc-genomics-workbench/) [58]. The expression values were expressed in reads per kilobase of exon model per million mapped reads (RPKM), and the normalized value for each sample was calculated in transcripts per million (TPM). For the analysis of differential expression, the following parameters were used: fold change greater than or equal to 1.5 and p-value lower than 0.05. The analysis of the exoproteome was performed by means of a BLASTn search of the predicted gene of ThIOC3844 against the local database of protein sequences from *T. harzianum* found in the extract of fungal growth under CEL and GLU conditions.

### Gene regulatory network

The gene regulatory networks were assembled from the reference mapped RNA-Seq data using each set of biological triplicates for the CEL and GLU conditions [35]. The interaction between the genes was obtained by calculating Pearson’s correlation for each pair of genes. The induction and repression networks were constructed based on the expression data of a set of genes that were identified in the secretome of the CEL growth condition by the Bayesian inference method [59]. If the secreted protein was present in the condition, it was assigned a value of one. If the secreted protein was absent, it was assigned a value of zero. The treatment conditions were considered as regulators of the network to detect the direct relationships between the conditions and the genes. Thus, the Bayesian network represents the relationships among the conditions, gene expression, and secreted proteins. Cytoscape software v 3.4.042 [60] (https://cytoscape.org/) was used for data analysis and construction of the CLR2 subnetwork.

## Additional files

**Additional file 1: Fig S1**. Screening genes of interest in the genomic library of *T. harzianum* IOC3844 by qPCR (a); reads size sequenced using PACBio technology (b); genes cluster in a genomic region of *T. harzianum* (c). **Fig. S2**. Distribution of the main GO terms of the annotated genes in *T. harzianum* IOC3844. **Fig. S3**. Pipeline approach for the analyzes used in this work of genes and genomic study in *T. harzianum*. **Supplementary Table S2**. Assembly parameters of a set of sequenced genomic region using PACBio technology. **Supplementary Table S3**. Comparison of genomic data among different species of *Trichoderma* spp. **Supplementary Table S8**. Description of the species used for the phylogenetic analysis of the transcription factor CLR2. **Supplementary Table S9**. Description of the genes found in the coregulation networks.

**Additional file 2: Supplementary Table S1**. Description of the genomic regions sequenced in *T. harzianum* IOC3844.

**Additional file 3: Supplementary Table S4**. Annotation of all genes predicted in *T. harzianum* IOC3844.

**Additional file 4: Supplementary Table S5**. Description of the EC codes for *T. harzianum* IOC3844 genes.

**Additional file 5: Supplementary Table S6**. Description of the CAZymes genes for *T. harzianum* IOC3844.

**Additional file 6: Supplementary Table S7**. Level of expression of the genes annotated in *T. harzianum* IOC3844 by means of RNA-seq.

## List of abbreviations

AA: Auxiliary enzymes;
B: buried;
BAC: bacterial artificial chromosome;
BLAST: Basic local alignment search tool;
bp: Base pair;
BRENDA: Braunschweig Enzyme Database;
C: loop;
CAZymes: Carbohydrate-active enzymes;
CBMAI: Brazilian Collection of Environment and Industrial Microorganisms;
CBM: Carbohydrate-binding module;
CE: Carbohydrate esterases;
CEL: Cellulose;
CIP1: cellulose-induced protein 1;
CIP2: cellulose-induced protein 2;
CLR2: cellulose degradation regulator 2;
DNA: Deoxyribonucleic acid;
E: beta-sheet;
EC: Enzyme commission number;
Ex: exposed;
FC: fold change;
GH: Glycoside hydrolases;
GLU: Glucose;
GO: gene ontologies;
GT: Glycosyltransferases;
H: helix;
JTT: Jones-Taylor-Thornton;
kb: Kilobases;
LPMO: Lytic polysaccharides monooxygenase;
M: medium;
Mb: Megabase;
MEGA: Molecular evolutionary genetics analysis;
MFS: major facilitator superfamily permease;
ML: maximum likelihood;
PacBio: Pacific Biosciences;
PBcR: PacBio Corrected Reads;
PCR: Polymerase chain reaction;
PL: Polysaccharide lyases;
RNA: Ribonucleic acid;
RNA-Seq: RNA sequencing;
RPKM: Reads per kilobase of exon model per million mapped reads;
SMRT: Single-Molecule Real-Time;
TaIMI206040: *T. atroviride* IMI206040;
TFs: transcription factors;
ThB97: *T. harzianum* B97;
ThIOC3844: *Trichoderma harzianum* IOC-3844;
ThTR274: *T. harzianum* TR274;
Th6766: *T. harzianum;*
TPM: Transcripts per million;
TrQM6a: *T. reesei* QM6a;
TvGv29-8: *T. virens* Gv29-8

## Declarations

### Ethics approval and consent to participate

Not applicable

### Consent for publication

Not applicable

### Availability of data and materials

The RNA-seq data can be accessed by the accession number <u>PRJNA336221</u>. Data from the genomic regions were submitted to GenBank (https://www.ncbi.nlm.nih.gov/genbank/) under the accession numbers MK861589-MK861650 (Supplementary Table S1).

### Authors ‘ contributions

APS and JAFF designed the study. JAFF, MACH, CAS, DAA, JSM, DAS and AC performed the research. JAFF, MACH, CAS, DAA, NFM and CBCS analyzed the data. JAFF, MACH, CAS and APS wrote the paper. All authors critically read the text and approved the manuscript.

## Acknowledgements

We would like to acknowledge the funding from Fundação de Amparo à Pesquisa do Estado de São Paulo (FAPESP 2015/09202-0), Coordenação de Aperfeiçoamento de Pessoal de Nível Superior (CAPES, Computational Biology Program) and Conselho Nacional de Desenvolvimento Científico e Tecnológico (CNPq).

