## Additional file 1 for "“Integrative Genomic Analysis for the Bioprospection of Regulators and Accessory Enzymes Associated with Cellulose Degradation in a Filamentous Fungus (*Trichoderma harzianum*)”"

**Supplementary data**

**This PDF file includes:**

Figs S1 to S3

Tables S2, S3, S8 and S9

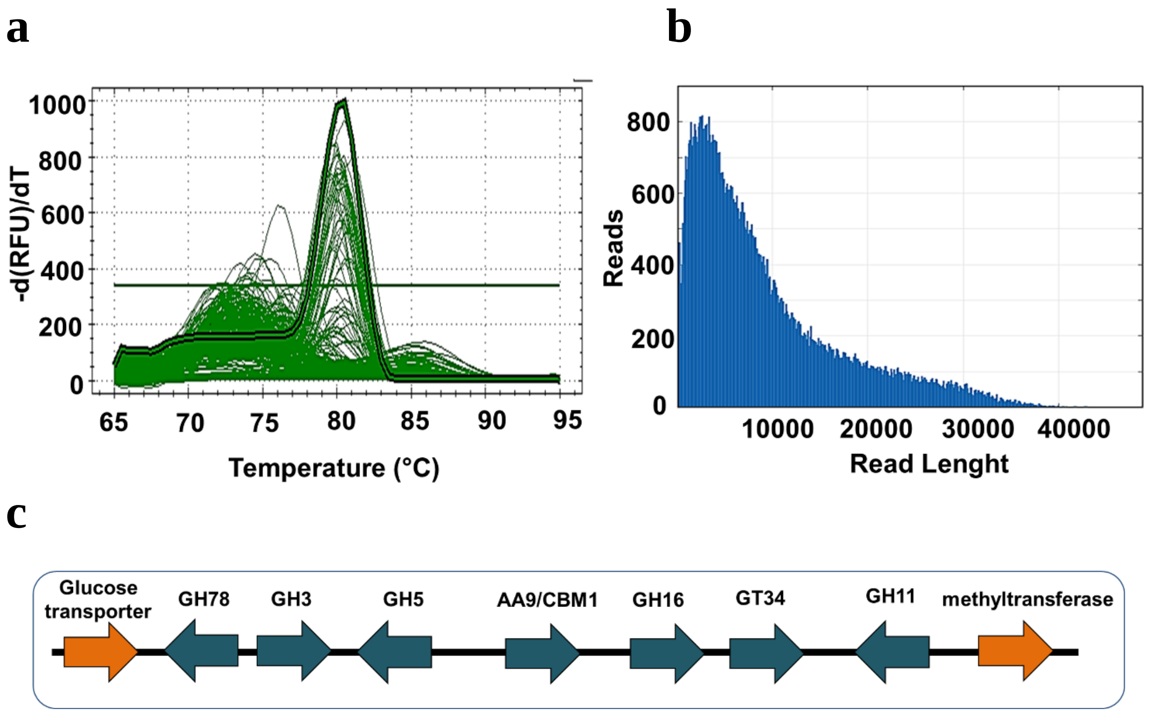


**Fig. S1.** Screening genes of interest in the genomic library of *T. harzianum* IOC3844 by qPCR (a); reads size sequenced using PACBio technology (b); genes cluster in a genomic region of *T. harzianum* (c).

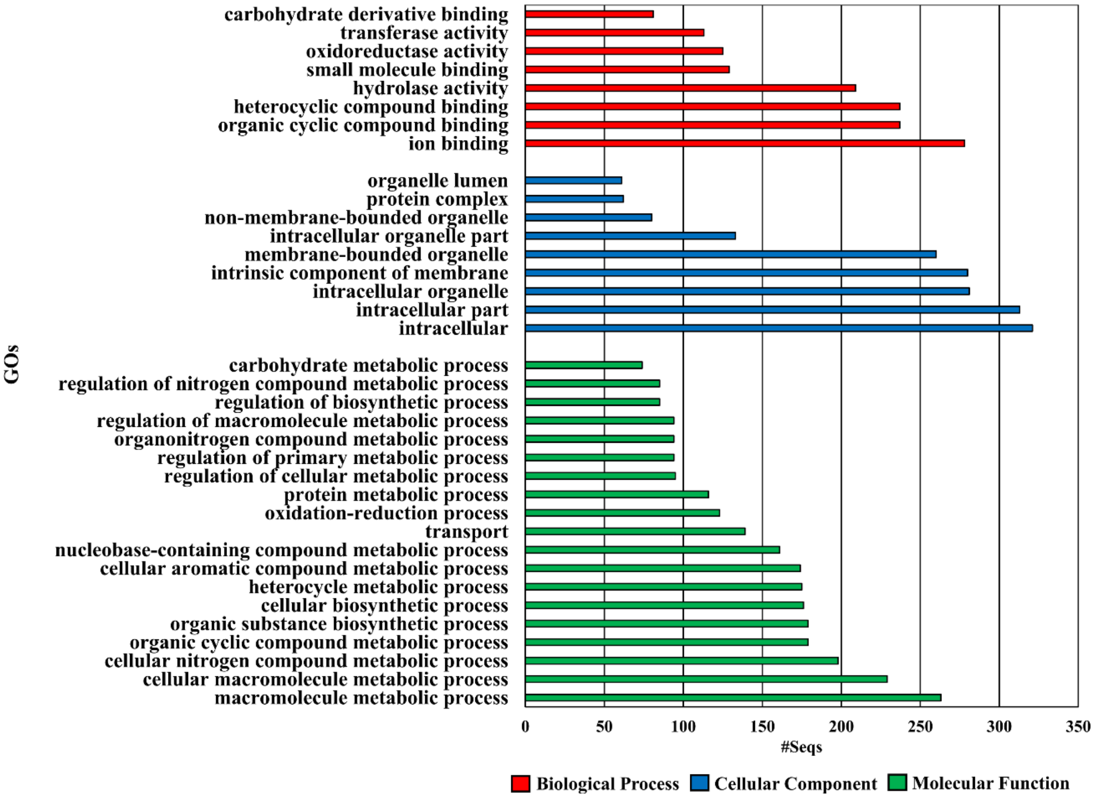


**Fig. S2.** Distribution of the main GO terms of the annotated genes in *T. harzianum* IOC3844.

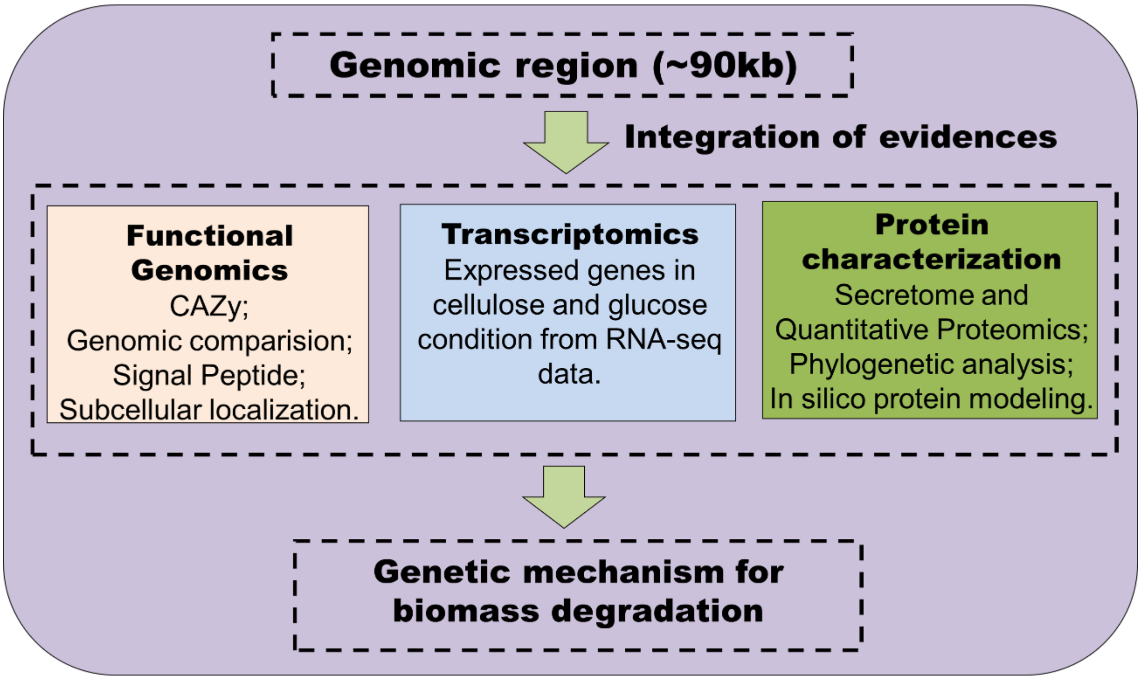


**Fig. S3.** Pipeline approach for the analyzes used in this work of genes and genomic study in *T. harzianum*.

**Supplementary Table S2.** Assembly parameters of a set of sequenced genomic region using PACBio technology.

| Parameters | Results |
|---|---|
| Polymerase Read Bases | 1151876133 |
| Seed Bases | 150009116 |
| Pre-Assembly Yield | 0.296 |
| Pre-Assembly Reads Length | 5528 |
| Length Cutoff | 12815 |
| Pre-Assembly bases | 44419992 |
| Pre-Assembly Reads | 8035 |
| Pre-Assembly N50 | 8199 |
| Mean Read Score | 0.667 |
| Number of Reads | 100732 |
| Mean Read Length | 11493 |
| Number of bases | 1157715225 |
| N50 Read Length | 17608 |
| Polymerase Read Quality | 0.849 |

**Supplementary Table S3.** Comparison of genomic data among different species of *Trichoderma* spp.

| Organism/Name | Strain | Genome type | Size (Mb) | GC% | Scaffolds | CAZymes |
|---|---|---|---|---|---|---|
| ***T. harzianum*** | IOC3844 | BACs | 5.00 | 49.65 | 63 | 222 |
| ***T. harzianum*** | T6776 | Draft | 40.98 | 48.50 | 1572 | 430 |
| ***T. reesei*** | QM6a | Complete genome | 33.39 | 52.80 | 7 | 327 |
| ***T. atroviride*** | IMI 206040 | Draft | 36.14 | 49.70 | 29 | 422 |
| ***T. virens*** | Gv29-8 | Draft | 39.00 | 49.20 | 93 | 441 |

**Supplementary Table S8.** Description of the species used for the phylogenetic analysis of the transcription factor CLR2.

| Group | Sequence ID | Species |
|---|---|---|
| 1 | XP_012182108.1 | *Fibroporia radiculosa* |
| | KZT72632.1 | *Daedalea quercina* L-15889 |
| | PCH41399.1 | *Wolfiporia cocos* MD-104 SS10 |
| | XP_024336022.1 | *Postia placenta* MAD-698-R-SB12 |
| | OCH94255.1 | *Obba rivulosa* |
| | PSR73669.1 | *Phlebia centrifuga* |
| | XP_007392271.1 | *Phanerochaete carnosa* HHB-10118 |
| | XP_008035817.0 | *Trametes versicolor* FP-101664 SS1 |
| | XP_007364283.1 | *Dichomitus squalens* LYAD-421 |
| | OJA15282.1 | *Rhizopogon vesiculosus* |
| | OAX40653.1 | *Rhizopogon vinicolor* AM-OR11-026 |
| | KIJ70318.1 | *Hydnomerulius pinastri* MD-312 |
| | KIK30790.1 | *Pisolithus microcarpus* 441 |
| | KIO13022.1 | *Pisolithus tinctorius* Marx 270 |
| 2 | OXG14340.1 | *Cryptococcus neoformans* var. grubii Tu2591 |
| | OWZ51650.1 | *Cryptococcus neoformans* var. grubii 125.91 |
| | OXG77158.1 | *Cryptococcus neoformans* var. grubii Br795 |
| | OWZ35064.1 | *Cryptococcus neoformans* var. grubii AD1-83a |
| | XP_012051901.1 | *Cryptococcus neoformans* var. grubii H99 |
| | OWT36835.1 | *Cryptococcus neoformans* var. grubii Bt1 |
| | OXG13205.1 | *Cryptococcus neoformans* var. grubii Tu401-1 |
| | OWZ76252.1 | *Cryptococcus neoformans* var. grubii Bt85 |
| | OXM75578.1 | *Cryptococcus neoformans* var. grubii Bt63 |
| | XP_018260593.1 | *Kwoniella dejecticola* CBS 10117 |
| | XP_018999125.1 | *Kwoniella mangroviensis* CBS 8507 |
| | XP_019045940.1 | *Kwoniella bestiolae* CBS 10118 |
| | XP_019014217.1 | *Kwoniella pini* CBS 10737 |
| | XP_018266380.1 | *Kwoniella dejecticola* CBS 10117 |
| | OCF60695.1 | *Kwoniella mangroviensis* CBS 10435 |
| | OCF74556.1 | *Kwoniella mangroviensis* CBS 8886 |
| | XP_019001153.1 | *Kwoniella mangroviensis* CBS 8507 |
| | XP_021869321.1 | *Kockovaella imperatae* |
| | XP_019045848.1 | *Kwoniella bestiolae* CBS 10118 |
| | XP_018260036.1 | *Kwoniella dejecticola* CBS 10117 |
| | XP_003196556.1 | *Cryptococcus gattii* WM276 |

| | | |
|---|---|---|
| 2 | OWZ29927.1 | *Cryptococcus neoformans* var. grubii c45 |
| | XP_019010711.1 | *Kwoniella pini* CBS 10737 |
| | POY75593.1 | *Rhodotorula taiwanensis* |
| | XP_018280147.1 | *Cutaneotrichosporon oleaginosum* |
| | EKD01798.1 | *Trichosporon asahii* var. asahii CBS 8904 |
| | XP_014181811.1 | *Trichosporon asahii* var. asahii CBS 2479 |
| 3 | CDM38126 | *Penicillium roqueforti* FM164 |
| | XP_660973.1 | *Aspergillus nidulans* FGSC A4 |
| 4 | PKK47514.1 | *Trichoderma harzianum* CL102_9848 |
| | Th_IOC3844 | *Trichoderma harzianum* IOC3844 |
| | XP_024778108.1 | *Trichoderma harzianum* CBS 226.95 |
| | KKP03054.1 | *Trichoderma harzianum* |
| | XP_013955330.1 | *Trichoderma virens* Gv29-8 |
| 5 | ETS03553.1 | *Trichoderma reesei* RUT C-30 |
| | XP_006964036.1 | *Trichoderma reesei* QM6a |
| | XP_013941191.1 | *Trichoderma atroviride* IMI 206040 |
| | XP_024764890.1 | *Trichoderma asperellum* CBS 433.97 |
| | KOS18048.1 | *Escovopsis weberi* |
| 6 | CZT51594.1 | *Rhynchosporium secalis* |
| | CZT06835.1 | *Rhynchosporium agropyri* |
| | CZT07141.1 | *Rhynchosporium commune* |
| | PMD16076.1 | *Pezoloma ericae* |
| | PMD30568.1 | *Hyaloscypha variabilis* F |
| | XP_024739234.1 | *Meliniomyces bicolor* E |
| | XP_018061285.1 | *Phialocephala scopiformis* |
| | CZR63207.1 | *Phialocephala subalpina* |
| | XP_007288749.1 | *Marssonina brunnea* |
| | PQE16296.1 | *Rutstroemia* sp. NJR-2017a WRK4 |
| | PQE03599.1 | *Rutstroemia* sp. NJR-2017a BBW |
| | ESZ95397.1 | *Sclerotinia borealis* F-4128 |
| | EMR81895.1 | *Botrytis cinerea* BcDW1 |
| | XP_024552100.1 | *Botrytis cinerea* B05.10 |
| | CCD51977.1 | *Botrytis cinerea* T4 |
| | KFX95591.1 | *Pseudogymnoascus* sp. VKM F-3557 |
| | OBT56849.1 | *Pseudogymnoascus* sp. 24MN13 |
| | KFZ12187.1 | *Pseudogymnoascus* sp. VKM F4519 (FW-2642) |
| | XP_018129406.1 | *Pseudogymnoascus verrucosus* |
| 7 | KUI68282.1 | *Valsa mali* |

| | | |
|---|---|---|
| 7 | KUI55118.1 | *Valsa mali* var. pyri |
| | POS76166.1 | *Diaporthe helianthi* |
| | KKY36004.1 | *Diaporthe ampelina* |
| | PSS02243.1 | *Coniella lustricola* |
| 8 | XP_016589835.1 | *Sporothrix schenckii* 1099-18 |
| | KIH94787.1 | *Sporothrix brasiliensis* 5110 |
| | EPE08882.1 | *Ophiostoma piceae* UAMH 11346 |
| 9 | XP_009849767.1 | *Neurospora tetrasperma* FGSC 2508 |
| | XP_962712.2 | *Neurospora crassa* OR74A |
| | XP_003347695.1 | *Sordaria macrospora* k-hell |
| | XP_006693821.1 | *Chaetomium thermophilum* |
| | KXX83256.1 | *Madurella mycetomatis* |
| | XP_003660436.1 | *Thermothelomyces thermophila* |
| 10 | KLU85969.1 | *Magnaporthiopsis poae* ATCC 64411 |
| | XP_009224916.1 | *Gaeumannomyces tritici* R3-111a-1 |
| | XP_003714853.1 | *Magnaporthe oryzae* 70-15 |
| 11 | OTA63138.1 | *Hypoxylon* sp. EC38 |
| | OTA97472.1 | *Hypoxylon* sp. CO27-5 |
| | OTB08885.1 | *Hypoxylon* sp. CI-4A |
| | OTB14150.1 | *Daldinia* sp. EC12 |
| | KXJ95218.1 | *Microdochium bolleyi* |
| 12 | ELA25990.1 | *Colletotrichum fructicola* Nara gc5 |
| | ENH84300.1 | *Colletotrichum orbiculare* MAFF 240422 |
| 13 | OLN97553.1 | *Colletotrichum chlorophyti* |
| 14 | KXH28512.1 | *Colletotrichum nymphaeae* SA-01 |
| | KXH27412.1 | *Colletotrichum simmondsii* |
| | XP_022478568.1 | *Colletotrichum orchidophilum* |
| | KXH59612.1 | *Colletotrichum salicis* |
| | XP_008097168.1 | *Colletotrichum graminicola* M1.001 |
| | KDN69204.1 | *Colletotrichum sublineola* |
| | KZL75422.1 | *Colletotrichum tofieldiae* |
| | OHW92794.1 | *Colletotrichum incanum* |
| | KZL81134.1 | *Colletotrichum incanum* |

**Supplementary Table S9.** Description of the genes found in the coregulation networks.

| Network ID | Protein ID | Protein name |
|---|---|---|
| 1 | KKP06817.1 | Hypotetical protein |
| 2 | KKP05702.1 | Hypotetical protein |
| 3 | KKP00617.1 | Hypotetical protein |
| 4 | KKP03936.1 | Kinase DC2 |
| 5 | KKP07737.1 | Translation initiation factor 3 subunit k |
| 6 | KKP01671.1 | Hypotetical protein |
| 7 | KKO98378.1 | Translation initiation factor subunit 1 |
| 8 | KKP03537.1 | Murein transglycosylase |
| 9 | KKO96961.1 | Hypotetical protein |
| 10 | KKP00812.1 | Hypotetical protein |
| 11 | KKP06268.1 | Hypotetical protein |
| 12 | KKP01476.1 | Peptidyl-prolyl-cis-trans isomerase sspl |
| 13 | KKO97887.1 | Hypotetical protein |
| 14 | KKO99717.1 | Hypotetical protein |
| 15 | KKP00810.1 | Hypotetical protein |
| 16 | KKP07167.1 | Hypotetical protein |
| 17 | KKP02416.1 | Hypotetical protein |
| 18 | KKO98063.1 | Vacuolar ATP synthase subunit D |
| 19 | KKP03148.1 | Hypotetical protein |
| 20 | KKP07719.1 | Hypotetical protein |
| 21 | KKO97889.1 | Hypotetical protein |
| 22 | KKP00888.1 | ATP-dependent RNA helicase DED1 |
| 23 | KKP02267.1 | Ubiquitin-protein ligase E3 C |
| 24 | KKP02972.1 | Hypotetical protein |
| 25 | KKO98726.1 | DUF718 |
| 26 | KKP02131.1 | Hypotetical protein |
| 27 | KKO96618.1 | Hypotetical protein |
| 28 | KKO98708.1 | Hypotetical protein |
| 29 | KKP02590.1 | Proteasome subunit alpha type-2 |
| 30 | KKO99933.1 | Hypotetical protein |
| 31 | KKP07598.1 | Undercaprenyl diphosphate synthase |
| 32 | KKP00804.1 | 14-3-3 family protein |
| 33 | KKP02280.1 | ATP-dependent RNA helicase SUB2 |
| 34 | KKP03054.1 | Cellulose Degradation regulator 2 – CLR2 |
| 35 | KKO97002.1 | Hydroxyacylglutathione hydrolase |
| 36 | KKP01112.1 | Hypotetical protein |
